# Neural pathways of voice perception: a comparative dMRI study between humans and macaques

**DOI:** 10.64898/2025.12.16.691583

**Authors:** Mélina Cordeau, Katherine L. Bryant, David Meunier, Régis Trapeau, Clémentine Bodin, Kep Kee Loh, Olivier Coulon, Pascal Belin

## Abstract

The processing of vocal stimuli in the human brain involves multiple regions that are selectively tuned to voice information. Similarly, voice-sensitive areas have been identified in macaques, suggesting a common neural mechanism across species. However, the structural connectivity underlying voice perception in both humans and macaques remains poorly understood. In this study, we investigate the structural connectivity of voice-sensitive regions in 4 humans and 3 macaques using awake functional MRI and diffusion MRI techniques. Our findings reveal that while humans and macaques differ in the anatomical organization of projections from the anterior Temporal Voice Area (aTVA), the organization of projections from the primary auditory cortex (A1) is nearly identical. Our results also demonstrate distinct connectivity patterns between the vocal areas of the two species, as well as an opposite pattern of lateralization of aTVA-A1 connectivity in humans and macaques. These results enhance our understanding of the neural circuits involved in voice perception and provide valuable insights into the comparative neurobiology of vocal communication.

## Introduction

The neural processing of voice information has been found to engage multiple regions in the human cortex that are selectively tuned to vocal stimuli. These regions include the Temporal Voice Areas (anterior TVA: aTVA, middle TVA: mTVA, posterior TVA: pTVA), located in the secondary auditory cortex bilaterally (Belin et al., 2000; Pernet et al., 2015), as well as three regions located bilaterally in the frontal cortex which we refer to as the Frontal Voice Areas (anterior FVA: aFVA, middle FVA: mFVA, posterior FVA: pFVA; (Aglieri et al., 2018)). The FVA shows considerable inter-individual variability that appears to be related to sulcal anatomy (Cordeau et al., 2023). Moreover, activations sensitive to conspecific vocalizations have also been observed in macaques (Perrodin et al., 2011; Petkov et al., 2008) and one of them, called anterior TVA (aTVA), has been demonstrated as functionally homologous between the two species (Bodin et al., 2021).

Significant progress has been made in understanding the structural connectivity and white matter pathways originating from the auditory cortex. Foundational studies have demonstrated the organization of projections from the auditory cortex to various cortical and subcortical regions, including the thalamus, prefrontal areas, and the parietal and temporal lobes (Hackett, 2011; Pandya, 1995; Romanski et al., 1999). These pathways form the basis of auditory processing and contribute to complex functions like speech and language perception in humans, as well as vocal communication in non-human primates. Specific projections, such as those within the superior temporal gyrus (STG) (Upadhyay et al., 2008) and inter-hemispheric connections (Steinmann et al., 2014), have been highlighted as essential components of the auditory-linguistic network. However, while significant attention has been given to language-related functions, less is known about the structural connectivity underlying voice perception, a crucial auditory function likely shared between humans and macaques. To date, no study has comprehensively investigated the anatomical pathways connecting voice-sensitive regions in these species.

In this study, we establish a link between voice-sensitive regions that have been identified using awake functional MRI in both humans and macaques, and their structural connectivity. To do this, we performed a comparative analysis of (i) the anatomical organization of the functional homologs of aTVA and primary auditory cortex (A1) in both species and (ii) the structural connectivity of voice-sensitive regions and the inter-connectivity of aTVA and A1 in n=4 humans and n=3 macaques.

We observed that in macaques, aTVA projects to prefrontal regions predominantly via the ventral pathway, while human aTVA projects to prefrontal regions predominantly via the dorsal pathway. While A1 projections appeared to be similar between the two species, we observed distinct patterns of structural connectivity in the vocal areas of humans and macaques. We show that the connectivity between aTVA and A1 appears to exhibit an opposite pattern of lateralization in our humans and macaque participants.

## Materials and methods

### Subjects

Humans. Four native French human speakers were scanned (females; 23-38 years old). Participants gave written informed consent and were paid for their participation.

Macaques. Three adult rhesus monkeys (*Macaca mulatta*) were scanned, two females (M2, M3) and 1 male (M1) of 4, 5 and 8 years of age and weighing between 4, 5 and 11 kg. Macaques were born at the Station de Primatologie in Rousset and at SILABE, Strasbourg University. Each animal was implanted with a custom-made MRI-compatible head-post under sterile surgical conditions. The animals recovered for several weeks before being acclimated to head restraint via positive reinforcement (juice rewards). All experimental procedures were in compliance with the National Institutes of Health’s Guide for the Care and Use of Laboratory Animals and approved by the Ethical board of Institut de <u>Neurosciences</u> de la Timone (ref 2016060618508941).

### Experimental protocol

The experimental protocol follows that described in (Bodin, Trapeau, et al., 2021), and the data used in this study were acquired during that study. Four main categories of sounds were used in the awake fMRI experiment: human voices, macaque vocalizations, marmoset vocalizations, and non-vocal sounds, each containing 24 stimuli, for a total of 96 sound stimuli. The set of stimuli used during training was different from the one used during scanning in order to minimize familiarization effects. Human voices contained both speech (sentence segments from the set of stimuli used in a previous study (Moerel et al., 2012a), n = 12), and non-speech (vocal affect bursts selected from the Montreal Affective Voices dataset (Belin et al., 2008); n = 12), equally distributed into positive (pleasure, laugh; n = 4), neutral (n = 4) and negative (angry, fear; n = 4) vocalizations. Macaque vocalizations, kindly provided by Marc Hauser (Hauser, 1991), included both positive (coos 25%, n = 6, grunts 25%, n = 6) and negative (aggressive calls 25%, n = 6, screams 25%, n = 6) calls. Marmoset vocalizations, kindly provided by Asif Ghazanfar (Ghazanfar & Liao, 2018), were divided into supposed positive (trill 25%, n = 6), neutral (phee 25%, n = 6, twitter 25%, n = 6) and negative (tsik 25%, n = 6) calls. These three primate call categories contained an equal number of female and male callers.

Non-vocal sounds included both natural (living 25%, n = 6, non-living 25%, n = 6) and artificial sounds (human actions 25%, n = 6, or not 25%, n = 6) from previous studies from our group (Belin et al., 2000; Capilla et al., 2013) or kindly provided by Christopher Petkov (Petkov et al., 2008) and Elia Formisano (Moerel et al., 2012b). Stimuli were adjusted in duration, resampled at 48828 Hz and normalized by root mean square amplitude. Finally, a 10-ms cosine ramp was applied to the onset and offset of all stimuli. During experiments, stimuli were delivered via MRI-compatible earphones (S14, SensiMetrics, USA) at a sound pressure level of approximately 85 dB.

Functional scanning was done using an event-related paradigm with clustered-sparse acquisitions. MION (Monocrystalline Iron Oxide Nanoparticles) was used for all sessions of monkeys. No contrast agent was used for human participants. Human subjects were instructed, and monkeys subjects were trained, to remain still and awake in the scanner during auditory passive listening sessions lasting approximately one and a half hours.

### MRI acquisition

Human and monkey participants were scanned using the same 3-Tesla scanner (Siemens Prisma). Human participants were scanned using a whole-head 64-channel receiver coil (Siemens) in a single session including one T1-weighted anatomical scan (TR = 2.3 s, TE = 2.9ms, flip angle: 9°, matrix size = 192 × 256 × 256; resolution 1 × 1 x 1 mm3), one diffusion anatomical scan (SE-EPI sequence in AP and PA, 1,8mm isotropic voxel resolution, TR/TE=3002/73,6 ms), and two functional runs (sequences were obtained from the Center for Magnetic Resonance Research (CMRR, University of Minnesota) multiband EPI sequences, version R016a. Multiband acceleration factor: 4, TR = 0.945 s, TE = 30ms, flip angle = 65°, matrix size = 210 × 210 × 140, resolution of 2.5x2.5x2.5 mm3). We acquired 1 session per subject (2 runs of 96 stimulus presentation each), 841 volumes/run. Structural and diffusion anatomical scans were acquired with a 16-channel coil (ScanMed, Omaha, NE) for M1 monkey, and with a whole-head 24-channel receiver coil (Takashima) for M2 and M3 monkey. A high-resolution T1-weighted anatomical volume was acquired under general anesthesia (for M2, M3: MP2RAGE sequence, TE = 3.2ms, TR = 5 s, flip angle = 4°, matrix size = 176 × 160 × 160, resolution 0.4 × 0.4 × 0.4 mm3; for M1: MPRAGE sequence, TE = 3.15ms, TR = 3.3 s, flip angle = 8°, matrix size: 192 × 192 × 144, resolution 0.4 × 0.4 × 0.4 mm3), and also a diffusion anatomical scan (SE-EPI sequence in AP and PA, 1mm isotropic resolution, TR/TE=3500/81.8ms, multiband factor 2, partial Fourier: 3/4, 64 b1000, 3 repetitions in AP and PA). MION functional volumes were acquired with an 8-channels surface coil (KU, Leuven) using EPI sequences (multiband acceleration factor: 2, TR = 0.955 s, TE = 19ms, flip angle = 65°, matrix size = 108 × 108 × 48, resolution 1.5 × 1.5 × 1.5mm3). A total of 6 sessions were acquired in M1 (24 runs of 96 stimulus presentations each), 19 sessions were acquired in M2 (79 runs of 96 stimulus presentations each) and 21 sessions in M3 (72 runs of 96 stimulus presentations each). Between 841 and up to 1400 volumes/run, depend on the movements during the acquisition. Before and after each MION run, data were collected to allow the calculation of T2∗ maps by acquiring 18 volumes at the 3 gradient echo times of 19.8, 61.2 and 102.5ms.

### MRI preprocessing

Functional MRI. The functional data were preprocessed and contrast maps were computed in the previous study (Bodin, Trapeau, et al., 2021). Preprocessing of the functional data included motion correction, spatial distortion reduction using field maps, inter-run registration and spatial smoothing (SPM12 (Ashburner et al., 2021)).

Diffusion MRI. For both human and macaque, diffusion MRIs were preprocessed using the Diffuse pipeline implemented in Brainvisa (Brun et al., 2019). It performed corrections for motion, eddy-current and susceptibility induced distortions (Topup + eddy), and bias. Probabilistic fiber tractography was computed using MRtrix. MRtrix produces a human and macaque whole-brain tractography of 100 million streamlines and is corrected at 30 million after SIFT.

### MRI data analysis

Functional MRI. General linear model estimates of responses to all sounds versus silence (all > silence) and to conspecific vocalizations versus non-conspecific vocalizations (CV > non-CV) were computed using fMRISTAT (Worsley et al., 2002).

Structural connectivity. Probabilistic fiber tractography was computed using MRtrix. MRTRIX produces a whole-brain tractogram and a seed-based ‘filtered tractogram’ (and an estimate of the pair-wise strength of a pathway between two areas (Behrens et al., 2007; Johansen-Berg & Behrens, 2009)). The seed-based ‘filtered tractogram’ is based on the location of the functional peak (t-values). For the study of the anatomical projections from aTVA the location of the functional peaks was determined by the conspecific vocalization *vs.* all other sounds (from Bodin et al., 2021) (see Appendix A4); and for A1, the location of the functional peaks was determined by the sound *vs.* silence contrast. For connectivity between the voice patches: in humans, they were identified as TVA and FVA, as described in Aglieri et al., 2018 and Bodin, Trapeau, et al., 2021; in monkeys, the aTVA and A1 regions were determined based on the descriptions in Bodin et al., while other temporal and frontal peaks were individually selected among the strongest based on their activation intensity (t-values). Sphere sizes for the anatomical projections study (tractography based on a single seed, exploring all projections across the entire brain) are 10mm diameter for humans and 3.5mm diameter for macaques, because the anatomical surface diameter ratio is 2.8 between the two species. For the voice areas connectivity (tractography based on multiple seeds representing functional activations to specifically examine the connections between them), the location of the functional peaks was determined by the MNI coordinates described in Aglieri et al., 2018, as the closest local maxima at the individual level, and represented as a sphere. Sphere sizes for the connectome study (connectivity between voice patches and aTVA-A1 connectivity), were 10mm for human and 6.25mm for macaques, as the functional diameter ratio is 1,6 between the two species (ROI volume was 297 mm3 (diameter of 7.5 mm) in humans and 64 mm3 (diameter of 4.5 mm) in monkeys) (see Appendix A.5).

Connectivity strength. Connectivity strength is characterized by streamline number, that is, the number of tracts from each seed that successfully reach the target (Eickhoff et al., 2010; Forstmann et al., 2010). Probabilistic tracking was done between the voice-sensitive area in the superior temporal sulcus/gyrus and voice-sensitive areas in the frontal lobe. This was done for both of the identified TVA (Temporal Voice Area) and FVA (Frontal Voice Area) coordinates. One connectivity measure was calculated per pair of regions (symmetrical connectivity matrix).

## Results

### Anatomical projections from functional homolog A1 and aTVA

#### Projections from the primary auditory cortex A1

We examined the projections originating from A1 in both species. In Figure 1, we present individual-level visualizations. In both humans and macaques, A1 appears to be connected with the occipital, parietal, and frontal regions. The number of fibers between individuals (humans and macaques) appears to be nearly identical, confirming that our choice of ROI size (10mm for humans, 3.5mm for macaques) was relevant such that results of further analyses can be meaningfully compared. The precise count of streamlines extracted from the A1 ROI for each individual can be found in the appendix A.1 table.

**Figure 1:**
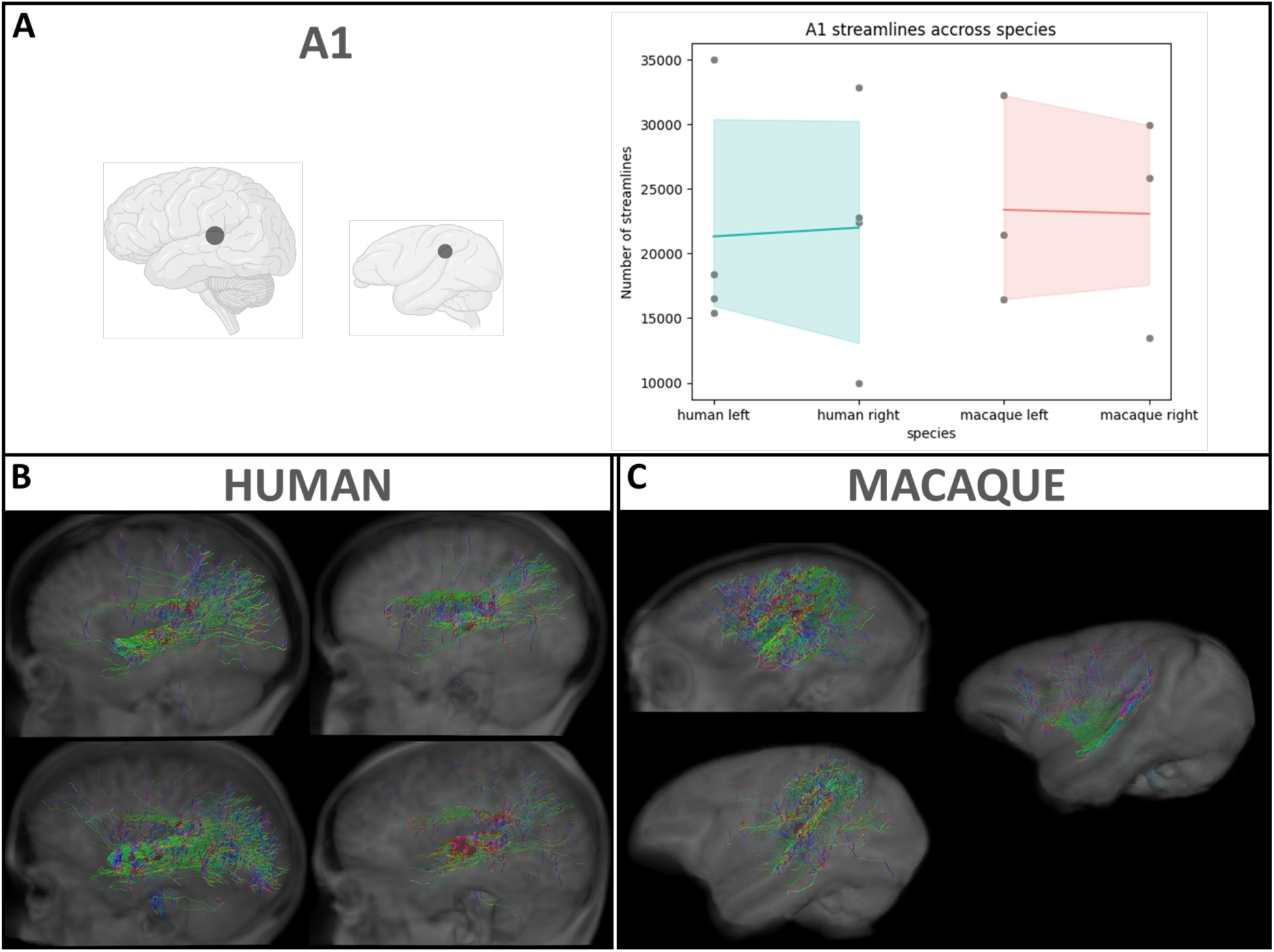
Individual-level identification of A1 (sound > silence) anatomical projections across the whole brain in our 4 humans and 3 macaques. A. The plot represents the number of streamlines originating from A1 for both species: humans in blue and macaques in pink. The thick line represents the fiber density for humans and macaques in the left and right hemispheres, while the blue and pink shading around it corresponds to the 95% confidence interval of the mean fiber density for each species. B and C. Visualization of anatomical projections from A1 ROIs (red spheres) for our 4 humans (sphere diameter reduced from 10mm to 5mm for visualization) and 3 macaques. Only the left hemisphere is shown. The colors in the tractography represent fiber directions: Green: Anterior-posterior, blue: Superior-inferior, red: Left-right.

#### Projections from aTVA

We studied the cortico-cortical projections originating from the functional activation aTVA in both humans and macaques. In Figure 2, we present individual-level visualizations. In humans, we observed that projections reach temporal, occipital, parietal, and frontal lobes. In macaques, projections are concentrated in the temporal and occipital lobes, and for one out of three individuals, in the frontal lobe. In addition to this difference, we observe that in humans, visually, the fibers predominantly seem to follow the dorsal pathway (fronto-temporal connections rather pass through the parietal lobe) rather than the ventral pathway, whereas in macaques, we do not observe a clearly dorsal or ventral pathway. In only one of the three macaques do we observe fibers projecting toward the frontal lobe, anterior to the temporal lobe. Furthermore, both monkeys had more projections originating from the right aTVA than the left aTVA. The precise count of streamlines extracted from the aTVA ROI for each individual can be found in the appendix A.2 table.

**Figure 2:**
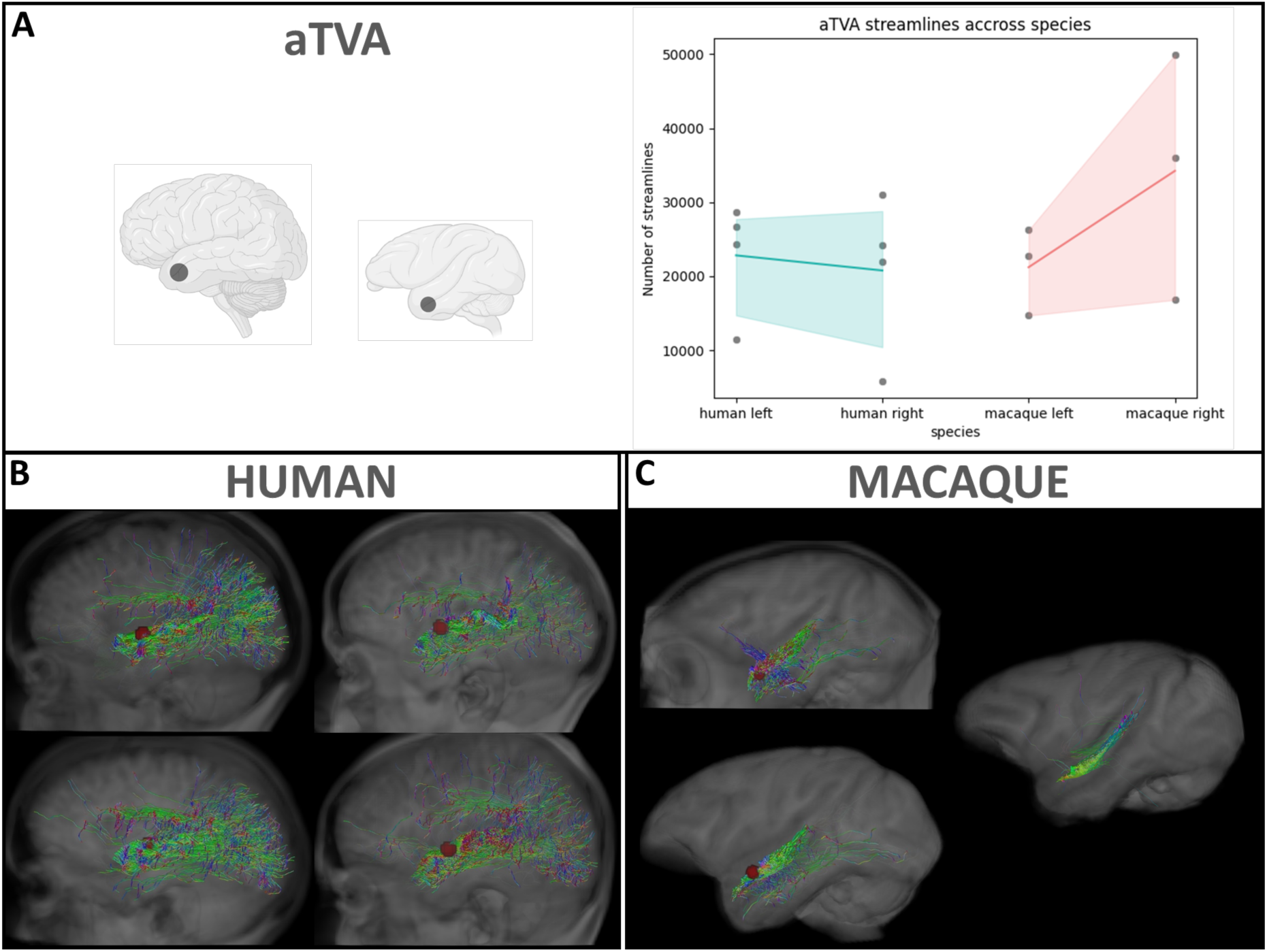
Individual-level identification of aTVA (conspecific > all other sounds) anatomical projections across the whole brain in our 4 humans and 3 macaques. A. The plot represents the number of streamlines originating from aTVA for both species: humans in blue and macaques in pink. The thick line represents the fiber density for humans and macaques in the left and right hemispheres, while the blue and pink shading around it corresponds to the 95% confidence interval of the mean fiber density for each species. B and C. Visualization of anatomical projections from aTVA ROIs (red spheres) for our 4 humans (sphere diameter reduced from 10mm to 5mm for visualization) and 3 macaques. Only the left hemisphere is shown. The colors in the tractography represent fiber directions: Green: Anterior-posterior, blue: Superior-inferior, red: Left-right.

### Structural connectivity of voice-sensitive regions in humans and macaques

#### Connectivity of voice patches

We investigated the structural connectivity of the voice areas in our 4 humans (TVAs and FVAs) and 3 macaques (aTVA, temporal activations and frontal activations). In Figure 3, we present visualizations of individual-level connections, the connectivity matrices. We observe that in humans, connectivity is noticeably stronger in the temporal lobe compared to the frontal lobe. Frontotemporal connections are present in humans but are very weak or even absent in macaques. We show that inter-hemispheric connections occur via the frontal areas in both species.

**Figure 3:**
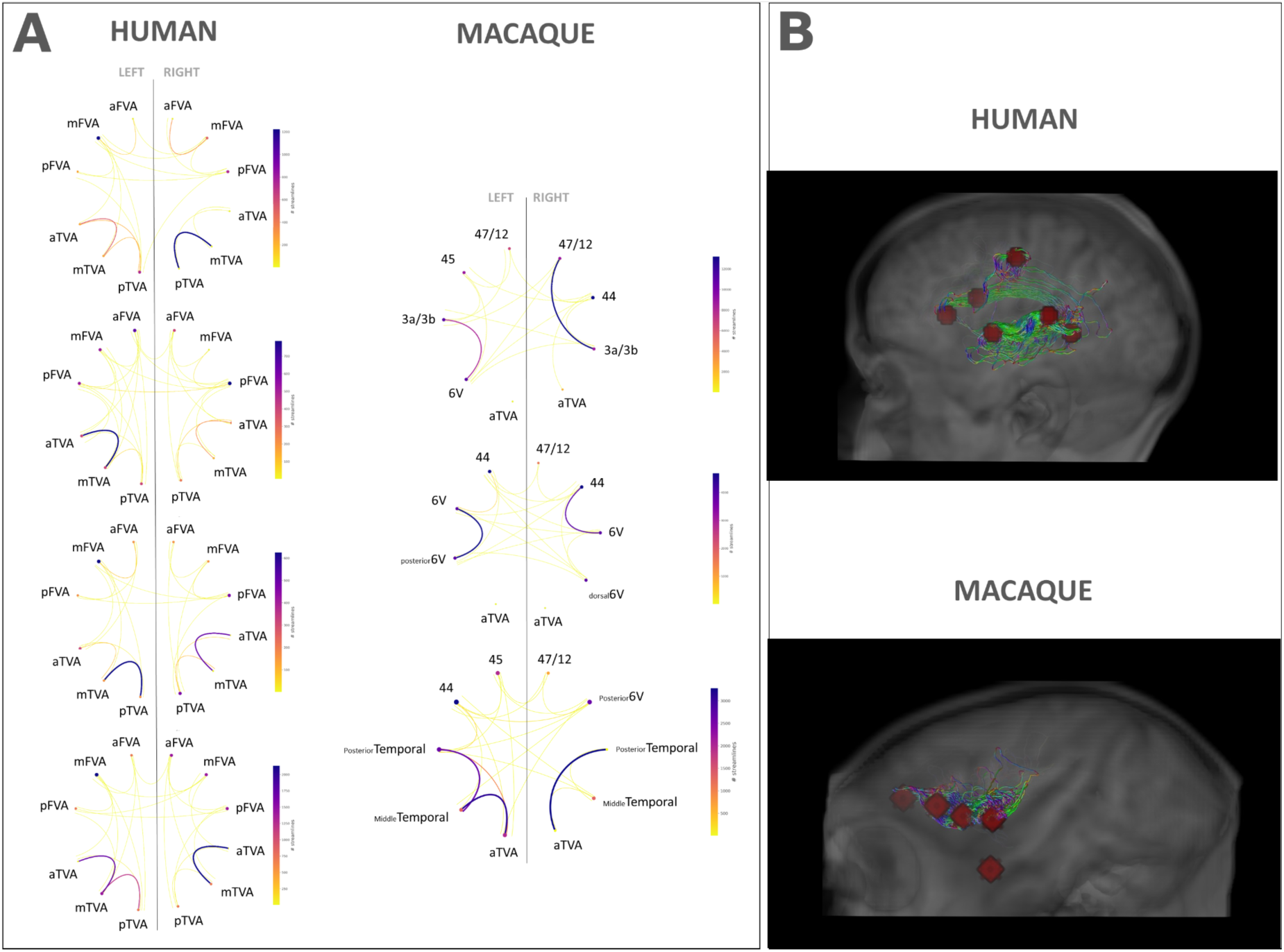
Visualizations of connectivity matrices in humans and macaques (A). In humans, matrices are comparable (3 TVA and 3 FVA for both hemispheres). In macaques, the 3 matrices are different because it depends on the location of the frontal activations. The matrices’ colorbar for both species represent the number of streamlines. On the right side we have one representative visualization per species (red spheres represent the individual functional activations) (B).

#### Connectivity between aTVA and A1

Next, we examined the connectivity between the functional homolog aTVA and the primary auditory cortex A1 for each individual. In Figure 4, we depict individual-level visualizations with quantifications of the number of connections between these two regions. In our humans, connectivity is much stronger in the left hemisphere (median: 672 streamlines; range: 828) than in the right hemisphere (median: 63 streamlines; range: 221). We observe a different pattern in our three macaques, with almost no direct connections in the left hemisphere (median: 3 streamlines; range: 8) and very few direct connections in the right hemisphere as well, except for one macaque who shows a large number of direct connections (median: 25 streamlines; range: 1950).

**Figure 4:**
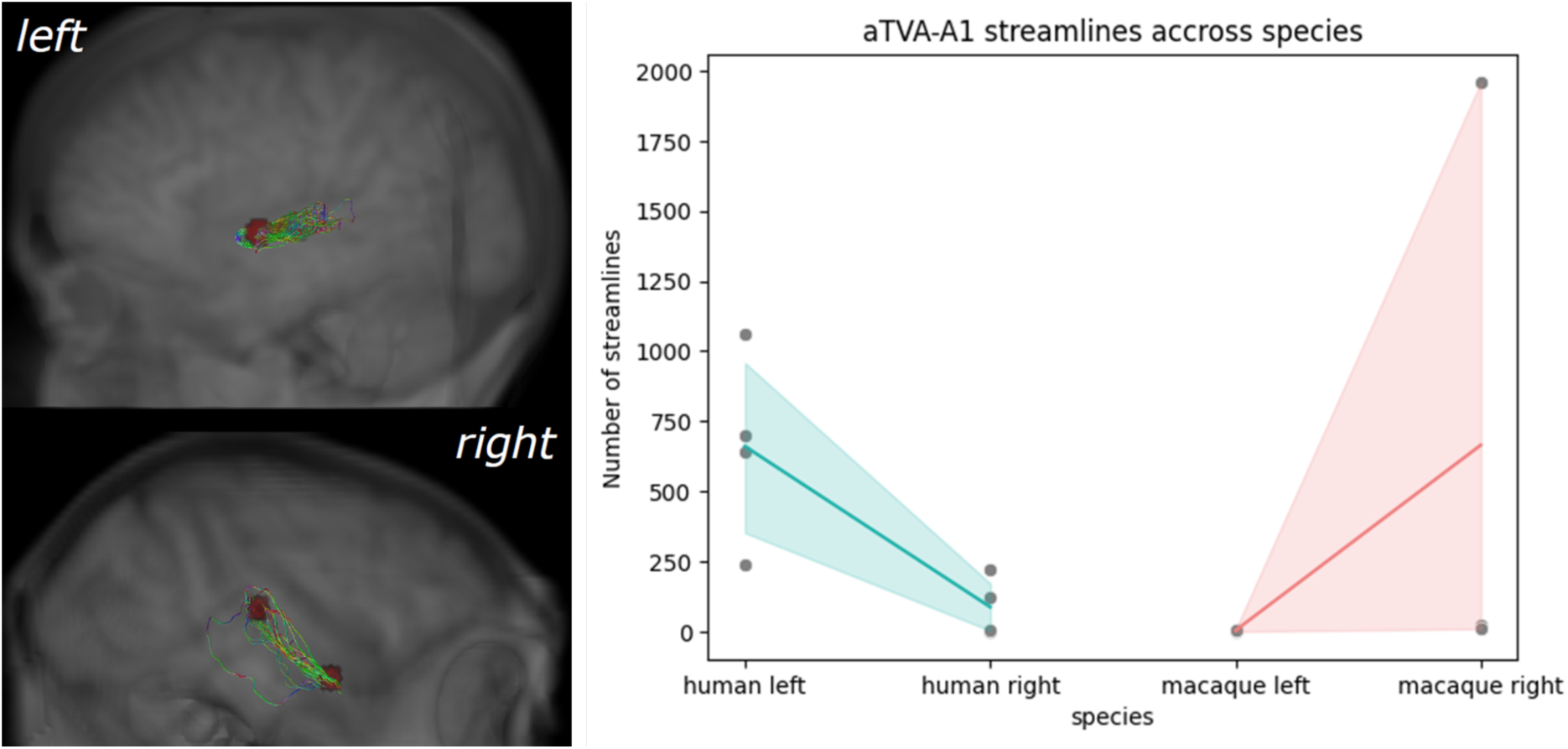
Visualizations of aTVA-A1 connectivity. A1 and aTVA are represented by the red spheres. Here we show an example of the connectivity in the left hemisphere of 1 human, and the right hemisphere of 1 macaque. This example allows us to highlight the reversed connectivity pattern that we observe in our 4 humans and 3 macaques. Plots represent the number of detected streamlines for each subject. Humans in green and macaques in red.

## Discussion

### A1 exhibits comparable anatomical organization in humans and macaques

Our investigation extended to an analysis of projections stemming from A1 activation in both humans and macaques. The visual representations at the individual level reveal remarkable patterns (fig. 2). In the context of humans, A1 connectivity extends to occipital, parietal, and frontal regions, primarily facilitated through the dorsal pathway (Hickok & Poeppel, 2007). Strikingly, a comparable connectivity pattern is observed in our macaques, where the primary auditory cortex, A1, also exhibits connections with the occipital, parietal, and frontal lobes. It was already shown that the primary auditory cortex is organized largely similarly in humans and macaques (Kaas & Hackett, 2000). Now, the robust presence of similar projections from A1 in both humans and macaques highlights potential shared underlying principles in auditory processing across these species. The concordance in projection patterns, despite evolutionary divergence, underscores the importance of these interconnected cortical pathways in auditory cognition.

### Comparative cortical projection patterns of aTVA in humans and macaques: unveiling anatomical connectivity and hemispheric specialization

The present study explores cortico-cortical projections stemming from the functional activation of aTVA in 4 humans and 3 macaques. We show distinct anatomical connectivity patterns associated with aTVA activation across these two species.

Concerning human subjects, our results described a robust distribution of projections originating from the area of activation in aTVA. These projections spanned multiple cortical regions, including the temporal, occipital, parietal, and frontal lobes. This extensive distribution suggests that aTVA activation in humans recruits diverse high-level cognitive functions. This observation aligns with previous research highlighting the multifarious roles of the temporal lobe in intricate cognitive processes such as language (Binding et al., 2022).

In contrast, the macaque functional homolog of aTVA displayed a more restricted projection pattern, primarily within the temporal and occipital lobes, with an additional projection in the frontal lobe for one of the three individuals. This species-specific variation in projection distribution may reflect evolutionary adaptations for species-specific cognitive functions.

Furthermore, an intriguing distinction emerged in terms of pathway prevalence between the two species. Human subjects exhibited a higher density of bundles in the dorsal pathway than in the ventral pathway, while macaques displayed the opposite trend (no bundle passed through the dorsal pathway, and one macaque showed projections to the frontal lobe). These results are in line with the study conducted by Balezeau and colleagues (Balezeau et al., 2020). The dorsal and ventral pathways are pivotal for auditory processing and sensory integration, respectively (Rauschecker & Tian, 2000; Romanski et al., 1999). The observed variations could arise from differences in auditory specialization and cognitive demands between humans and macaques.

The aTVA is recognized as a functional homolog in both humans and macaques (Bodin et al., 2021). The concept of homology contends that similar neural structures can serve comparable functions across different species (Striedter, 2005). However, the observation of a higher density of fiber bundles resulting from aTVA activation in the right hemisphere of macaques raises questions not only about hemispheric asymmetry but also about the potential preservation of this lateralized projection pattern across species. Hemispheric lateralization, often associated with functions like language processing, may also involve specializations in the right hemisphere (Vigneau et al., 2011), which could contribute to the formation of the observed right-lateralized connectivity. The shared functional homology of aTVA between humans and macaques encourages further investigations to determine if, beyond functional homology, similar cognitive processes are at play in both species.

In conclusion, the identification of a homologous functional role of aTVA, coupled with our observations of the underlying structural connectivity, calls for a more in-depth exploration of auditory processing mechanisms across species and the origin and evolution of vocal perception specialization.

### Distinct structural connectivity patterns in the Voice Areas of humans and macaques

We examined the structural connectivity of voice areas in both humans and macaques, in order to describe the neural pathways underlying auditory processing. The visual representations and connectivity matrices presented in Figure 3 provide a comprehensive view of the individual-level connections implicated in this pathway.

Notably, our analysis reveals a robust pattern of connectivity that is more pronounced within the temporal lobe when compared to the frontal lobe in humans. The strong temporal connectivity aligns with previous research highlighting the pivotal role of these regions in auditory processing and voice recognition (Belin et al., 2000) as well as their structural connectivity (Blank et al., 2011). We notice that in humans, the dorsal pathway (tracts running from the posterior temporal lobe to the frontal lobe) predominates over the ventral pathway (tracts running from the anterior temporal lobe to the prefrontal lobe), which is consistent with the study by (Balezeau et al., 2020). Additionally, we observe a connection between the aTVA and aFVA that we can presume to be either the UNC (Uncinate Fasciculus) or the IFOF (Inferior Fronto-Occipital Fasciculus). We cannot definitively determine which of these two bundles is actually involved in this connection, but we can hypothesize that it is the IFOF. Indeed, in the macaque, for this same fronto-temporal connection via the ventral pathway, we do not observe the UNC. However, we do observe, in one of the three macaques, a connection via the EmC (Extreme Capsule), a component of the IFOF (Mars et al., 2016). Therefore, we can imagine that the connection observed in the macaque belongs to the IFOF, and that this connection, or bundle, has evolved over time. This is also supported by studies by (Barrett et al., 2020; Bryant et al., 2020; Roumazeilles et al., 2020). This disparity in connectivity between species suggests potential species-specific adaptations or unique cognitive demands related to conspecific vocalization processing and communication systems.

Our investigation also sheds light on the inter-hemispheric connectivity patterns within these auditory circuits. Both humans and macaques exhibit inter-hemispheric connections, with the frontal areas serving as a conduit for this communication. This finding aligns with the notion of inter-hemispheric anatomical connections of the auditory cortex (Steinmann et al., 2014).

### Different connectivity pattern between aTVA-A1 in humans and macaques

Connectivity between the aTVA (anterior voice area) and A1 (primary auditory cortex) is essential to understanding the anatomical foundations of voice processing in humans and macaques. Our analysis of individual-level connections in both species reveals differences in connectivity patterns (Fig. 4). In humans, connectivity between aTVA and A1 shows a higher fiber density than in macaques. This finding is consistent with previous research showing that, as brain size increases, connectivity between multimodal areas intensifies, promoting the development of densely interconnected modules and, in parallel, long-range projections between these modules (Schmidt & Polleux, 2021).

A particularly intriguing result is the pattern of lateralization in aTVA–A1 connectivity observed between the two species. In humans, this pathway is left-lateralized, in line with the well-established role of the left hemisphere in language processing and auditory integration (Galaburda et al., 1978; Scott, 2005). In contrast, macaques showed no clear lateralization pattern, except for one individual who exhibited a right-hemisphere bias. This leftward lateralization in humans and the absence of lateralization—or presence of rightward lateralization in one macaque—suggest differences in the organization of auditory and potentially vocal functions across primate species.

It is important to place these findings within the broader context that brain structure and function lateralization is a characteristic feature of the primate lineage (Hecht et al., 2015; Hopkins, 2013; Sierpowska et al., 2022). This does not imply a fixed left- or right-hemisphere dominance, but rather that primate brains have the capacity to evolve lateralized organization in response to selective pressures—particularly for highly specialized behaviors such as vocal learning, language, or fine motor skills.

Based on the well-documented association between structural and functional lateralization in humans (Powell et al., 2006), we explored whether similar patterns would emerge in macaques for a vocal-related behavior. Since this is a vocal rather than a manual behavior, it was uncertain how lateralization would manifest in macaques. If they had shown left-hemisphere lateralization similar to humans, it would have suggested that the underlying trait evolved in a common ancestor. Instead, we observed a lack of lateralization overall, and an unexpected mirror image in one macaque subject. This opens two possible scenarios:

1. humans and macaques independently evolved structurally lateralized adaptations for vocal behavior;
2. primate brains have a general tendency to lateralize certain circuits (possibly due to plasticity during a critical developmental window), but the direction of this lateralization may vary across species or individuals.

The second scenario is supported by a growing number of recent review studies suggesting that brain lateralization in animals can result from genetic, environmental, epigenetic, or combined influences (Güntürkün et al., 2020), indicating that divergent developmental trajectories—such as species-specific environmental pressures—can lead to plasticity-driven lateralization patterns.

It is also worth noting that not all humans exhibit left-lateralized language processing—some individuals show right-hemisphere dominance (Knecht et al., 2000)—highlighting that lateralization is not absolute, but variable. This makes our interspecies findings even more compelling. Future studies with larger macaque samples will be essential to determine whether the absence of lateralization or the right-lateralization observed here is representative at the population level. Such data remain challenging to obtain, but this study represents a crucial first step toward identifying the evolutionary trajectories of lateralized voice processing networks in primates.

## Limitations

Several limitations of our study should be considered. First, there is no definitive guideline for choosing the number of streamlines generated for the whole-brain tractogram in humans and other primates. According to MRtrix recommendations, larger sampling yields better results. However, the optimal sampling for achieving a good distribution of seeds at the gray/white matter interface is not well-established. Furthermore, it is uncertain whether the same sampling should apply to different species, such as the macaque in our case. We opted to generate a tractogram with the same number of streamlines for both species, i.e., 100 million streamlines, as exemplified in BATMAN tutorial in MRtrix (Tahedl, 2020). We applied the SIFT algorithm to refine the tractograms, reducing them to 30 million streamlines. This ensures the streamline density better reflects the underlying fiber density from diffusion data. Given the smaller size of the macaque brain compared to that of humans, we acknowledge that we may have oversampled the macaque dataset. However, we aimed to mitigate this oversampling by reducing the size of the ROIs.

Regarding the choice of ROI size, we selected a 10mm diameter for humans based on visual assessment at the individual level and technical considerations for streamline filtering. Subsequently, we employed two different ROI sizes for macaques: for anatomical analyses (projections originating from a single ROI), we used a size ratio proportional to the gray/white matter interface surface to account for surface oversampling; for functional activation connectivity analyses, we employed a ratio proportional to the size of the activations.

A limitation in the study of A1-aTVA connectivity is that this result reflects the structural connectivity between two regions that are very close to each other. This means that the tractography method can be biased when it comes to short tracts. Therefore, we cannot determine whether the observed connections are direct or if we are obtaining a “smoothing” effect from a multitude of short-distance connections between these two regions (Saleem et al., 2000) or plis de passages (Bodin, Pron, et al., 2021).

## Conclusions

In conclusion, our study highlights both similarities and differences in the structural connectivity of the voice perception area homolog, aTVA, between humans and macaques. These findings support further investigation into auditory processing mechanisms and the evolution of specialized vocal perception.

Shared projections from A1 in both species suggest the existence of fundamental principles of auditory processing, while distinct frontotemporal connections in humans — and their presence or absence depending on individual macaques — underscore the importance of anatomical organization in functional specialization. Our results reveal species-specific similarities and differences, hemispheric asymmetries that would require statistical validation with a larger sample, as well as the impact of evolutionary adaptations on cortical circuits, enriching our understanding of vocalization processing across species.

## Statements and Declarations

The study is reported in accordance with the ARRIVE guidelines. All methods and experimental protocols were carried out in accordance with relevant guidelines and regulations. The datasets used and/or analysed during the current study available from the corresponding author on reasonable request.

## Funding

This work was funded by Fondation pour la Recherche Médicale (<u>AJE201214</u> to Pascal Belin and FDT202204015255 to Mélina Cordeau); Agence Nationale de la Recherche grants <u>ANR-16-CE37-0011-01</u> (PRIMAVOICE), <u>ANR-16-CONV-0002</u> (Institute for Language, Communication and the Brain), and <u>ANR-11-LABX-0036</u> (Brain and Language Research Institute); the Excellence Initiative of Aix-Marseille University (A∗MIDEX); and the European Research Council (ERC) under the European Union’s Horizon 2020 research and innovation program (grant agreement no. <u>788240</u>).

## Competing Interests

The authors have no relevant financial or non-financial interests to disclose.

## Author Contributions

MRI data were acquired by CB, RT and PB. fMRI data were preprocessed by CB, RT, and dMRI data by MC. fMRI data were analyzed by RT. Functional peak extraction was performed by MC, DM and KL. dMRI analysis was performed by MC and OC. Scripts and statistical analysis were performed by DM and MC. The manuscript was written by MC. All authors read and approved the final draft.

## Acknowledgement

This work was performed in the Center IRM-INT (UMR 7289, AMU-CNRS), platform member of France Life Imaging network (grant ANR-11-INBS-0006).

## Notes

### Competing Interest Statement

The authors have declared no competing interest.

