## Supplementary for "Neural pathways of voice perception: a comparative dMRI study between humans and macaques"

### Appendices

#### Appendix A.1 table: Number of detected streamlines originating from the A1 ROI

The number of counted streamlines originates from ROIs centered on the individual functional peak of A1 (sound *vs.* silence), and displaced to the nearest gray/white matter interface. The ROIs have a diameter of 10mm for humans and 3.5mm in macaques, as we utilized the gray/white matter surface size ratio between the two species.

| **A1** | **Number of streamlines Left** | **Nb streamlines Right** |
|---|---|---|
| H1 | 18368 | 32867 |
| H2 | 35013 | 22409 |
| H3 | 16521 | 22766 |
| H4 | 15393 | 9975 |
| M1 | 32255 | 29925 |
| M2 | 16471 | 13470 |
| M3 | 21425 | 25833 |

Human left: 17444 ± 19620 Human right: 22588 ± 22892

Macaques left: 21425 ± 15784 Macaques right: 25833 ± 16455

#### Appendix A.2 table: Number of detected streamlines originating from the aTVA ROI

The number of counted streamlines originates from ROIs centered on the individual functional peak of aTVA (conspecific vocal sounds *vs.* all other sounds), and displaced to the nearest gray/white matter interface. The ROIs have a diameter of 10mm for humans and 3.5mm in macaques, as we utilized the gray/white matter surface size ratio between the two species.

| **aTVA** | **Number of streamlines Left** | **Number of streamlines Right** |
|---|---|---|
| H1 | 26729 | 24192 |
| H2 | 28674 | 31039 |
| H3 | 11485 | 5794 |
| H4 | 24325 | 22005 |
| M1 | 26218 | 49929 |
| M2 | 14675 | 16798 |
| M3 | 22704 | 35948 |

Human left: 25527 ± 17189 Human right: 23098 ± 25245

Macaques left: 22704 ± 11543 Macaques right: 35948 ± 33131

#### Appendix A.3 table: Number of detected streamlines between A1 and aTVA ROIs

The number of counted streamlines between the functional activation for A1 and aTVA in each individual, and displaced to the nearest gray/white matter interface. The ROIs have a diameter of 10mm for humans and 6.25mm in macaques, as we utilized the functional activation size ratio between the two species.

| **A1-aTVA** | **Number of streamlines Left** | **Number of streamlines Right** |
|---|---|---|
| H1 | 641 | 1 |
| H2 | 236 | 121 |
| H3 | 1064 | 6 |
| H4 | 702 | 222 |
| M1 | 8 | 1960 |
| M2 | 0 | 25 |
| M3 | 3 | 10 |

Human left: 672 ± 828 Human right: 64 ± 221

Macaques left: 3 ± 8 Macaques right: 25 ± 1950

Appendix A.4 figure: Visualization of functional homologs aTVA in macaques, as previously defined in the paper by Bodin, Trapeau et al.

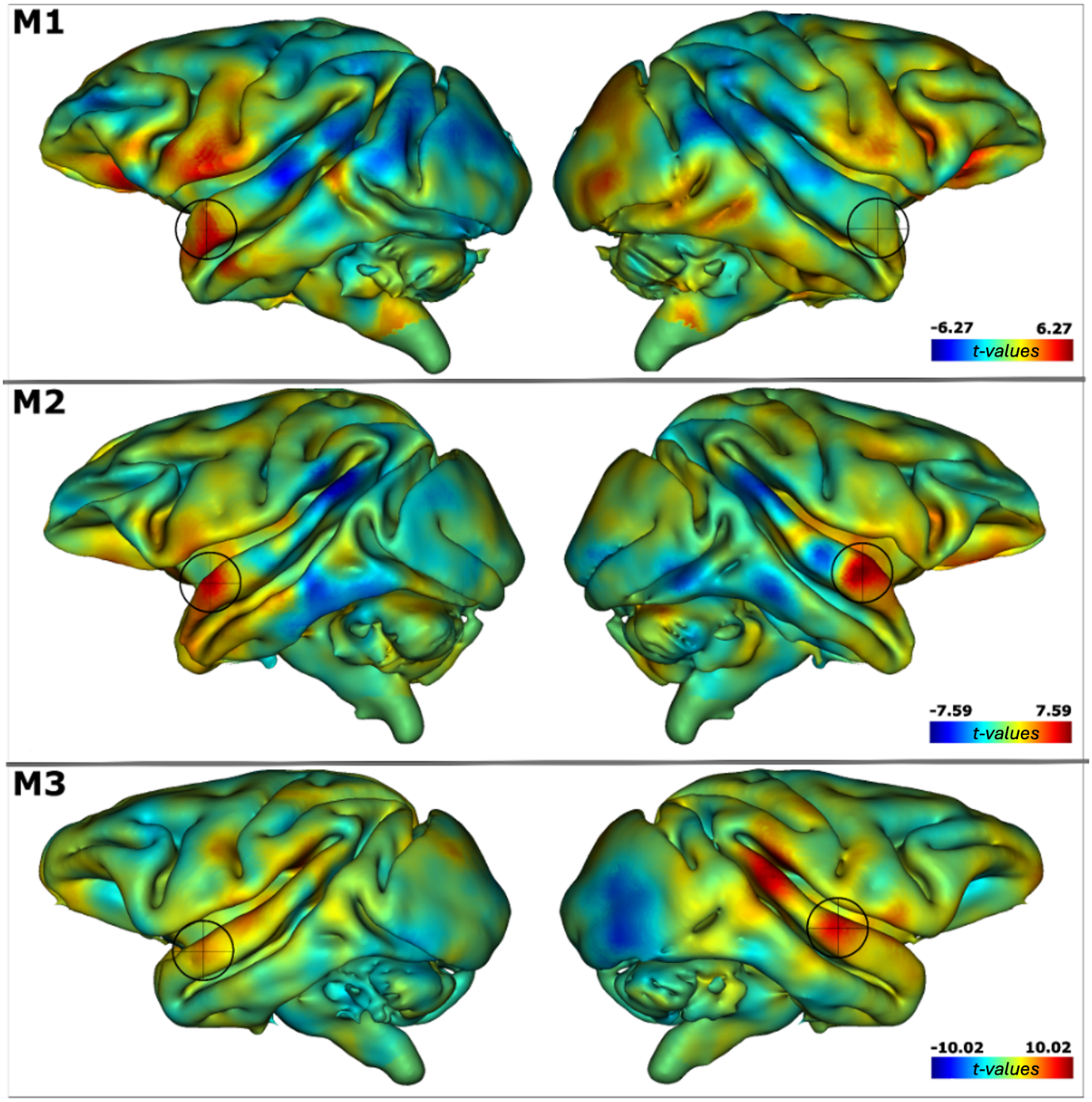

Appendix A.5 figure: Schematic representation of the location and size of the ROIs for each study and each species.

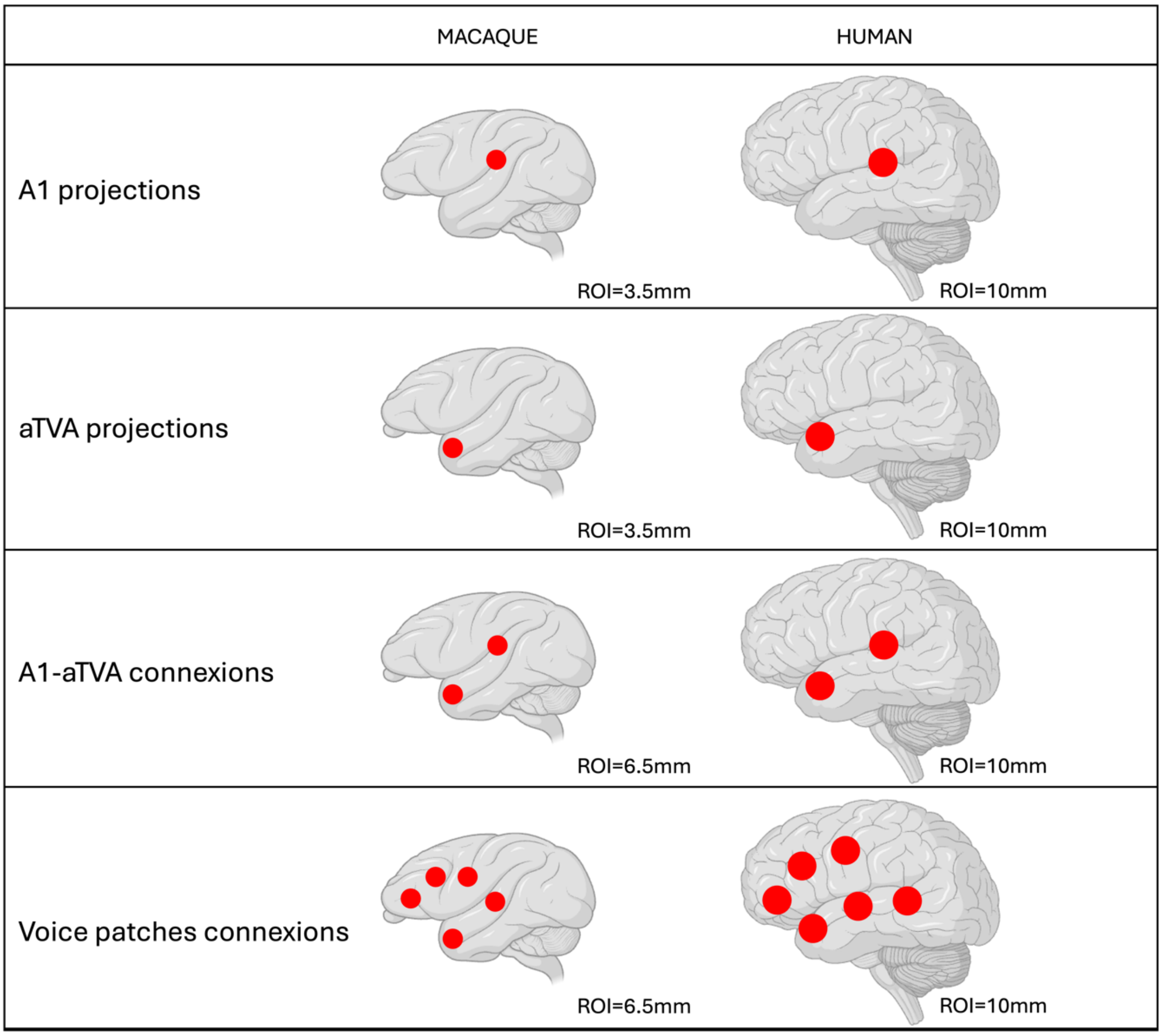

Appendix A.6 figure: Screenshot of the projections and connections observed at the individual level.

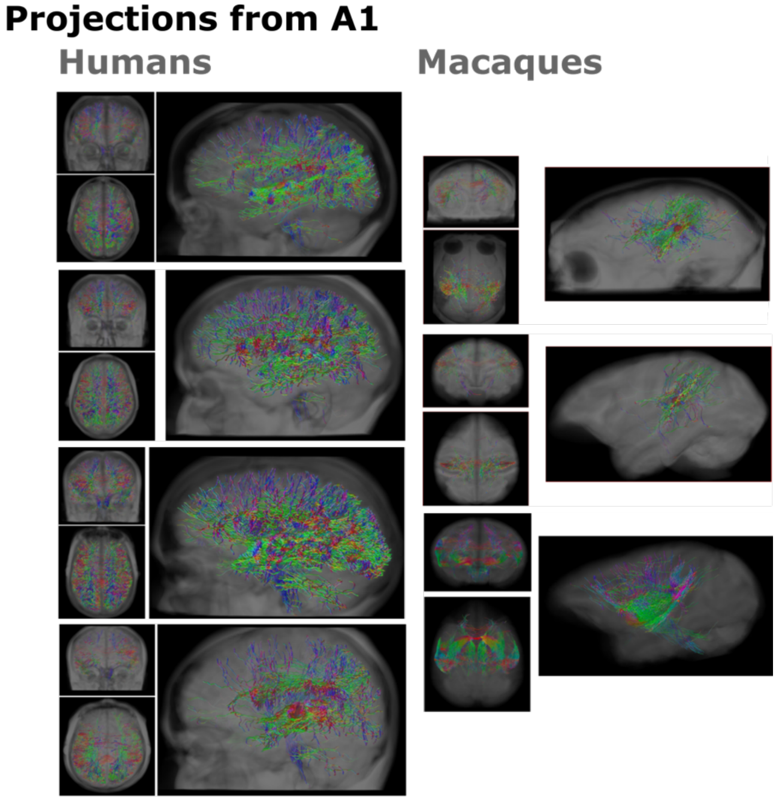


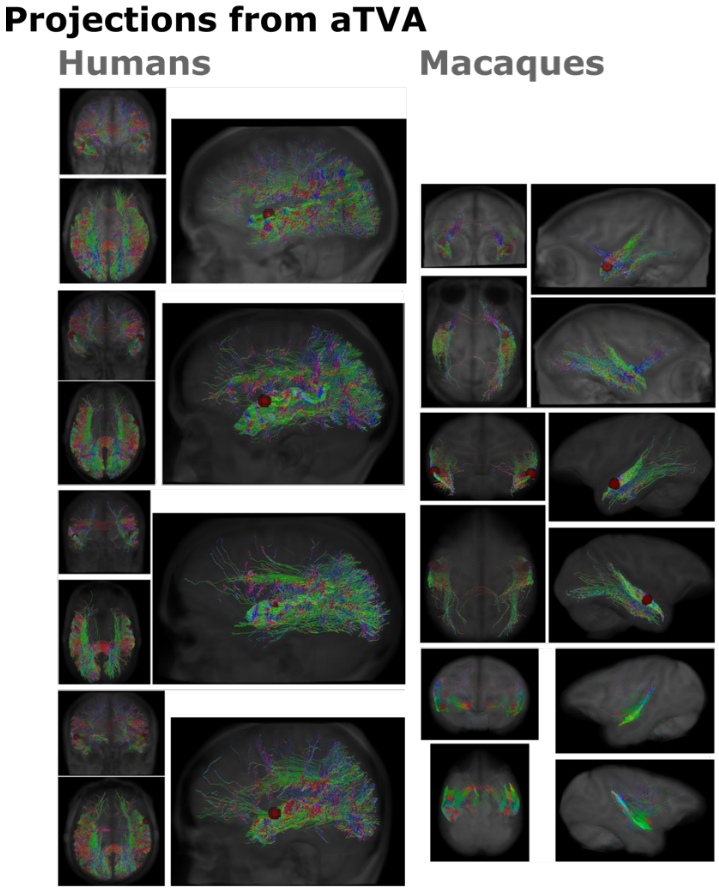

#### Connections between aTVA-A1

##### Macaques

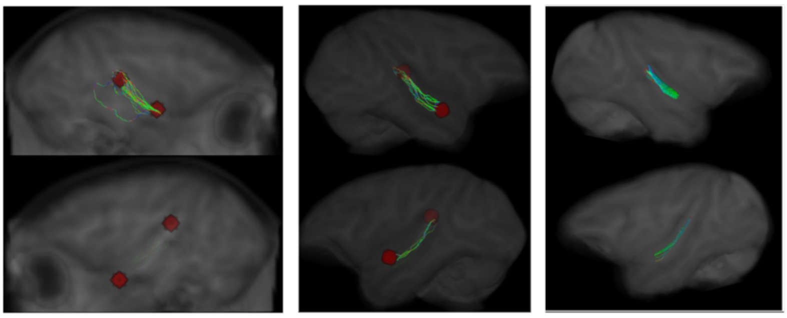

#### Connections between voice patches

##### Humans Macaques

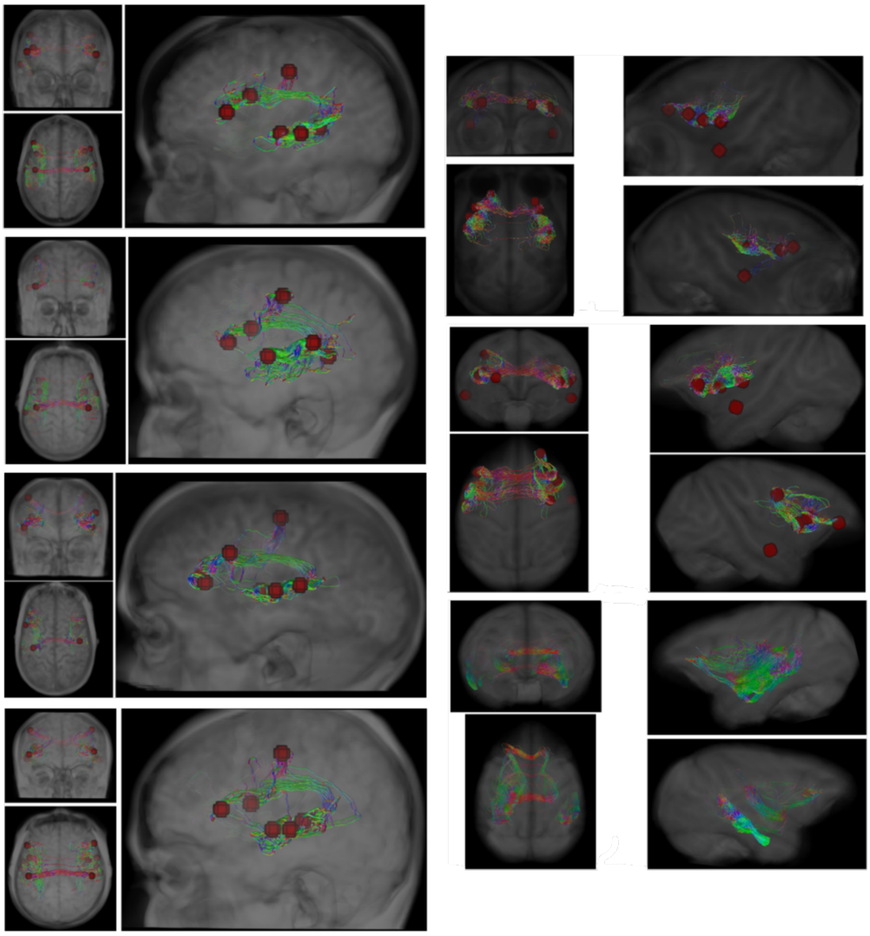
